# Reduced Lytic Activity of Bacteriophages in Presence of Antibiotics Targeting Bacterial Protein Synthesis

**DOI:** 10.1101/2021.06.30.450658

**Authors:** Medhavi Vashisth, Shikha Yashveer, Nitin Virmani, Bidhan Chandra Bera, Rajesh Kumar Vaid, Taruna Anand

## Abstract

Combination therapy of bacteriophage and antibiotics offers promise to treat multiple drug resistant bacterial infections through phage antibiotic synergy. However, its usage requires careful assessment as most antibiotics with mechanisms dependent upon inhibiting cell growth through interfering bacterial protein synthesis machinery were found to have an antagonistic effect on phage activity.

## Text

**V**arious alternatives to antibiotics including antimicrobial peptides, probiotics, antibodies and phytochemicals are being explored post emergence of antimicrobial resistance (1). Also, there is a substantial decrease in novel antibiotics being discovered and produced. Amongst the alternative antibacterial therapies, the usage of bacteriophages is an important consideration as they can infect bacteria and lyse them naturally. Several phage based therapeutic products are available and being used especially in Eastern Europe (2). Further, the usage of bacteriophages and antibiotics is in vogue as together they are known to enhance killing of bacteria by a phenomenon known as phage antibiotic synergy (PAS) (3). Recently some human cases of phage therapy have harnessed PAS for successful treatment of infections (4,5). However, the protocols to employ the bacteriophages and/or their combination with antibiotics for treatment of the infection in patients are not well elaborated and at times not substantiated scientifically (6). This makes it crucial to explore the antibiotics which show synergy with phages selected for the therapy.

Our team while exploring for PAS for two different bacteriophages (VTCCBPA145 and VTCCBPA182) (Figure 1) against *Acinetobacter baumannii* stumbled on an unusual phenomenon of antagonism with a selection of antibiotics instead. In brief, the screening for PAS was routinely planned in our laboratory for an *A. baumannii* isolate which is one of the rapidly emerging pathogens in the health care settings and leads to bacteremia, meningitis, urinary tract infections, pneumonia and wound infections (7). The bacteriophages in question VTCCBPA145 and 182 were being tested for assessment of synergy with different antibiotics by measuring the plaque size in the subminimal concentration zone surrounding the antibiotic disk as per the protocol of Comeau et al., 2008 (3). The antibiotics (13 nos.) screened against *A. baumannii* as per CLSI, guidelines included - Amikacin (AK), Gentamicin (GEN), Tobramycin (TOB), Ceftazidime (CAZ), Cefepime (CPM), Cefotaxime (CTX), Meropenem (MRP), Imipenem (IPM), Colistin (CL), Polymixin B (PB), Piperacillin (PI), Tetracycline (TE) and Ciprofloxacin (CIP). Out of these, CL and PI showed significant synergy (p<0.001) with both phages; CAZ, CPM, CTX, MRP and PB showed synergy with VTCCBPA182 but no effect on BPA145 was observed whereas GEN, TOB and TE showed significant (p<0.0001) antagonistic effect on both the phages. Intrigued with the finding we looked deeper into all aspects of antibiotics and noticed that only those antibiotics which act by inhibiting the cell growth through interfering with the bacterial protein synthesis machinery showed antagonistic effect with both bacteriophages.

**Figure 1:**
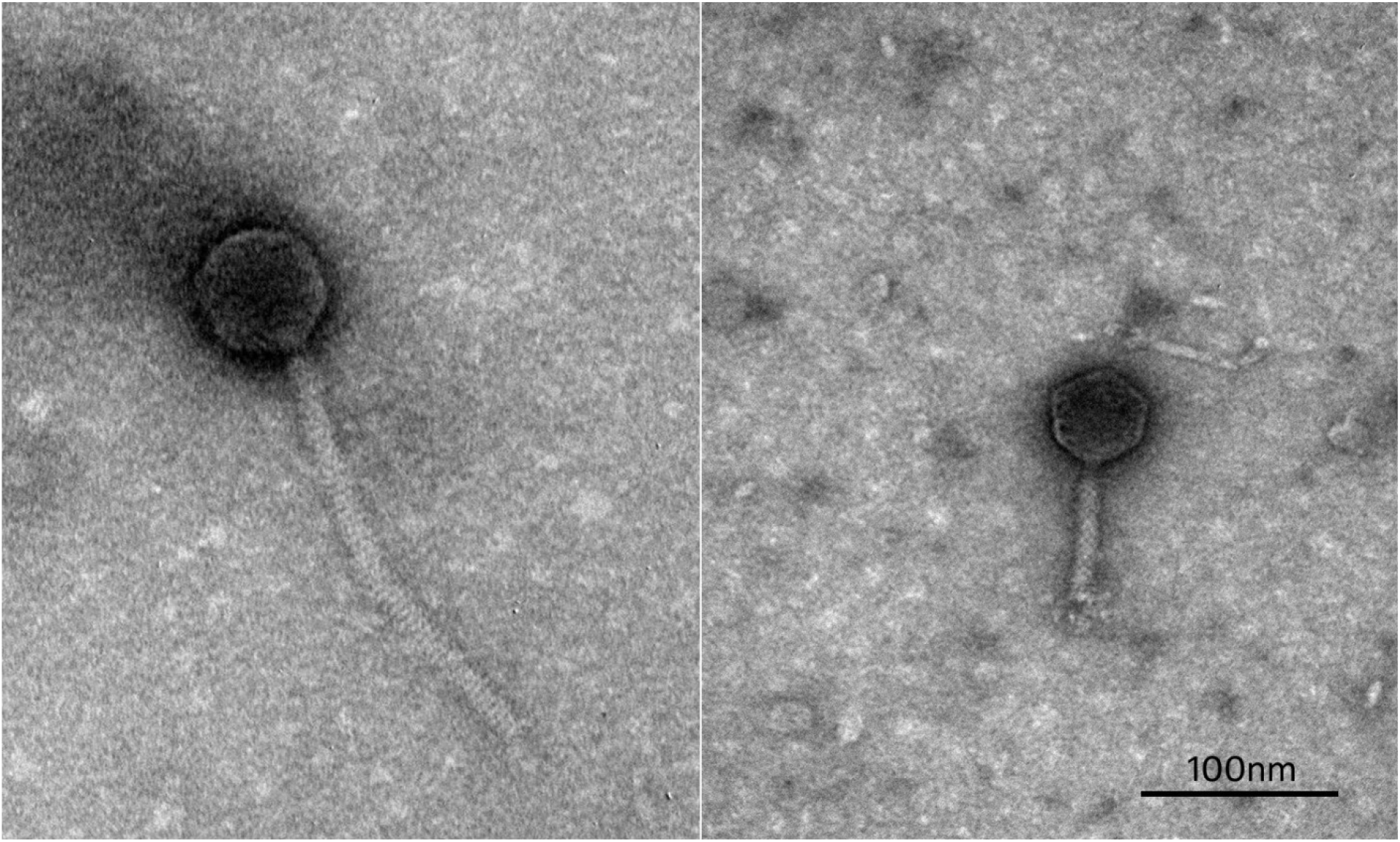
Transmission Electron Microscopy of BPA145 from family *siphoviridae* (left) and BPA182 from family *myoviridae* (right) using JEOL JEM-1011, Jeol, USA. Prior to visualization the bacteriophage suspensions were stained with 2% uranylacetete (pH4) on carbon coated nickel grids. Scale represents 100nm.

To confirm the antagonistic action of these antibiotics on phage lytic activity, 13 different antibiotics targeting the protein biosynthetic machinery [Amikacin (AK), Gentamicin (GEN), Kanamycin (K), Netilmicin (NET), Spectinomycin (SPT), Tobramycin (TOB), Tetracycline (TE), Oxytetracycline (O), Azithromycin (AZM), Erythromycin (E), Clindamycin (CD), Chloramphenicol (C) and Linezolid (LZ)] alongwith antibiotics targeting cell wall [Pipeacillin (PI), Cefepime (CPM) and Cefepime (CEP)] and cell membrane [Colistin (CL), Polymixin (PB)] of bacteria were further tested with a total of six bacteriophages. Two phages each against *Staphylococcus aureus* (VTCCBPA115 and VTCCBPA116), *Salmonella* Typhimurium (VTCCBPA143 and VTCCBPA188) and *A. baumannii* (VTCCBPA145 and VTCCBPA182) were used in the study. All the bacteriophages showed selective synergy with antibiotics targeting either cell wall or cell membrane but no antagonistic effect was observed with any such antibiotic. However bacteriophages showed highly significant antagonism in terms of decreased plaque size with most of the protein machinery targeting antibiotics: BPA115(13/13); BPA116 (13/13); BPA143(11/13); BPA145(11/13); BPA188(9/13) and BPA182(7/13) (Figure 2,3). We further confirmed the findings in liquid infection assays using *A. baumannii* bacteriophages *viz*. BPA145 and BPA182 in presence of sub-minimal inhibitory concentrations of GEN and TE. In both the cases, the antibiotic supplementation posed a significant antagonistic effect on bacteriophage lytic activity confirming further the proof of concept (Figure 4).

**Figure 2:**
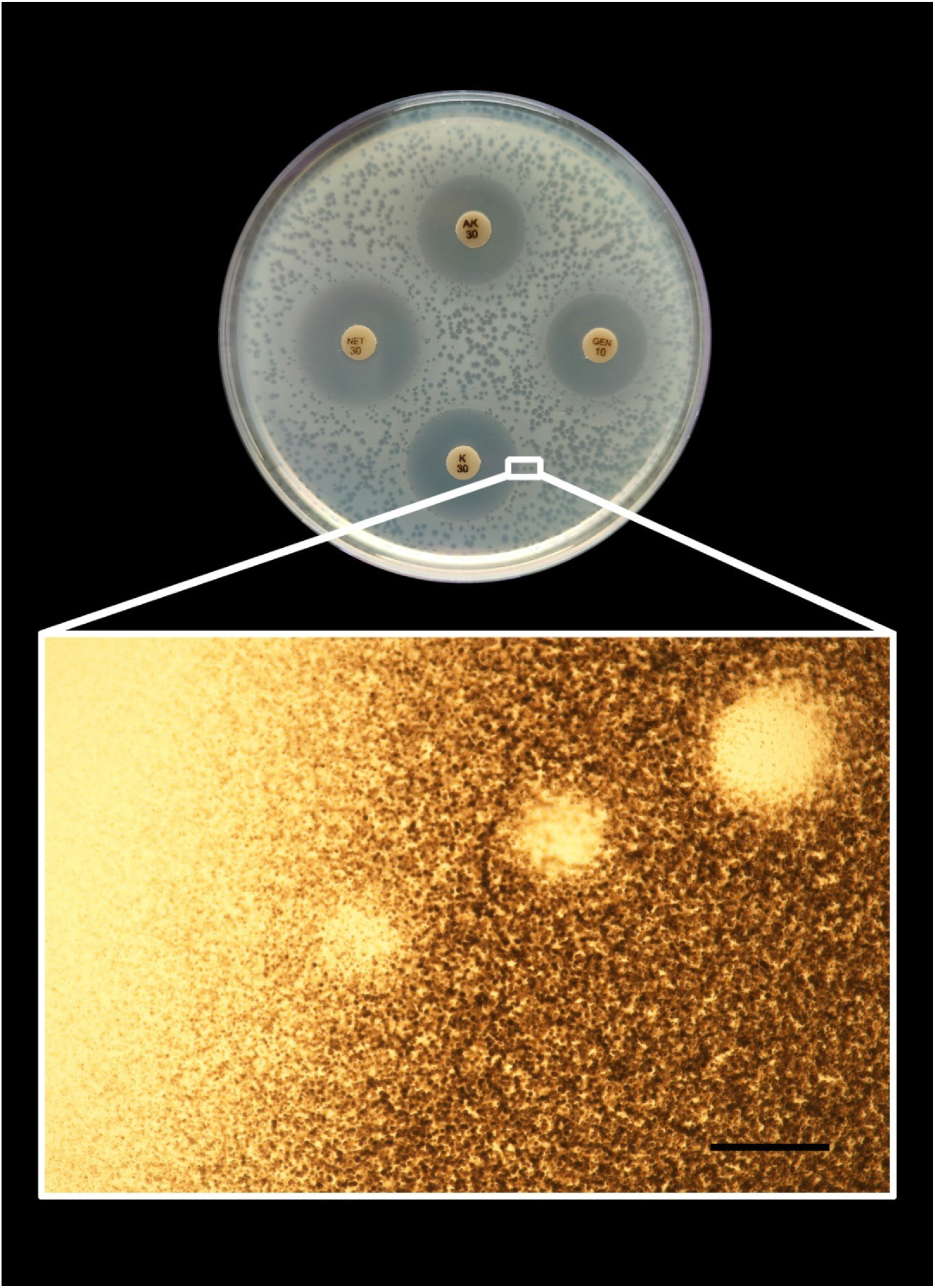
Plaque characteristics of bacteriophage BPA145 on double layer agar. The plaques in the vicinity of antibiotic disk inhibited bacterial zone have a smaller size as compared to the farther ones. Zoomed in picture shows the increase in plaque size with increased distance from the zone of clearance produced by antibiotic disk (40X magnification). Scale represents 500μm.

**Figure 3:**
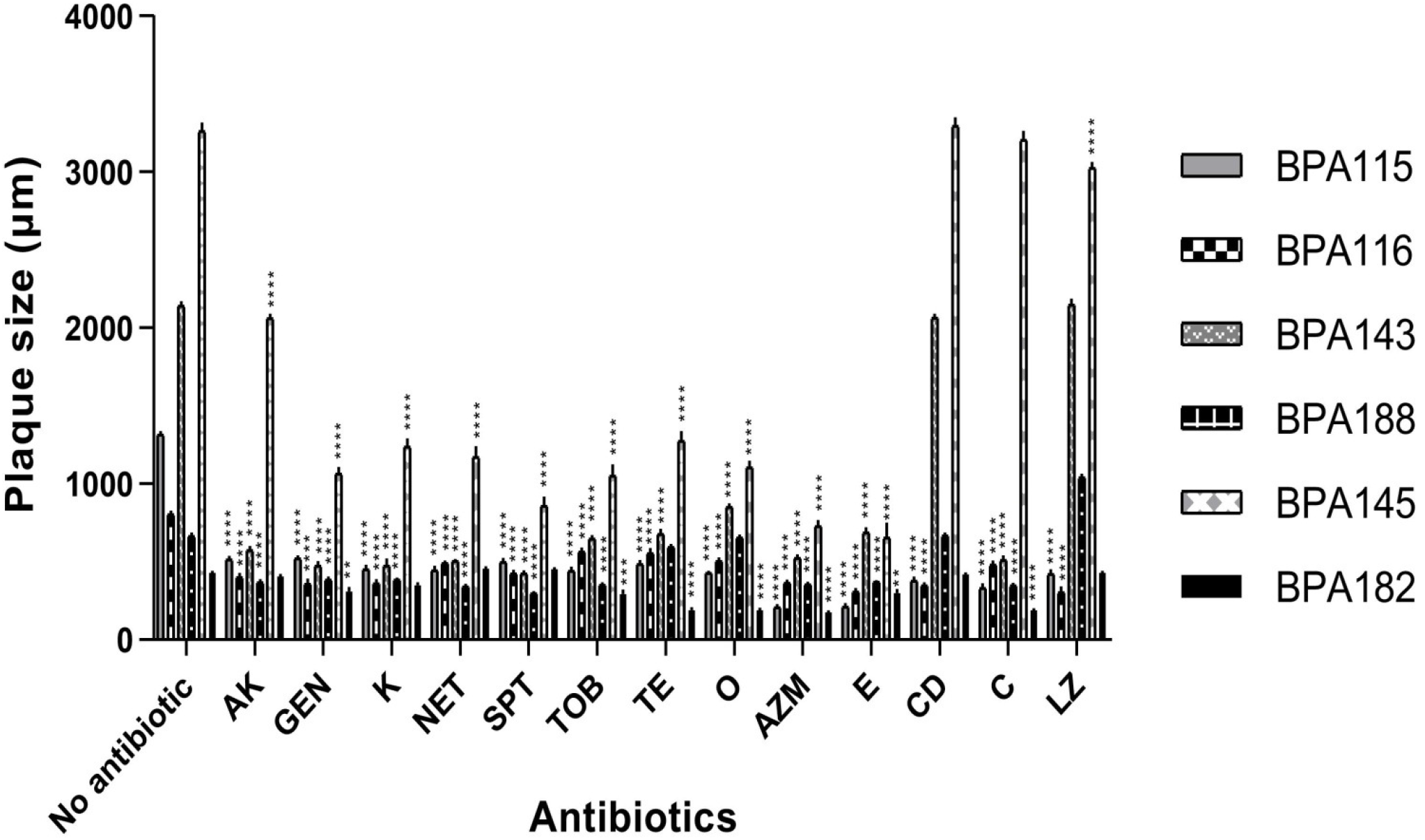
Change in plaque diameter as observed with different antibiotics measured in the vicinity of antibiotic inhibited bacterial zone. The plaque size (10nos.) was measured under 40X magnification in Nikon Eclipse Ni-E upright microscope. Statistical analysis was performed by two-way factorial analysis of variance using Dunnett's multiple comparisons test in GraphPad Prism 8.0.2. Significance levels are depicted as**** p<0.0001; *** p<0.001; ** p<0.01. Antibiotics are abbreviated as Amikacin (AK), Gentamicin (GEN), Kanamycin (K), Netilmicin (NET), Spectinomycin (SPT), Tobramycin (TOB), Tetracycline (TE), Oxytetracycline (O), Azithromycin (AZM), Erythromycin (E), Clindamycin (CD), Chloramphenicol (C) and Linezolid (LZ).

**Figure 4:**
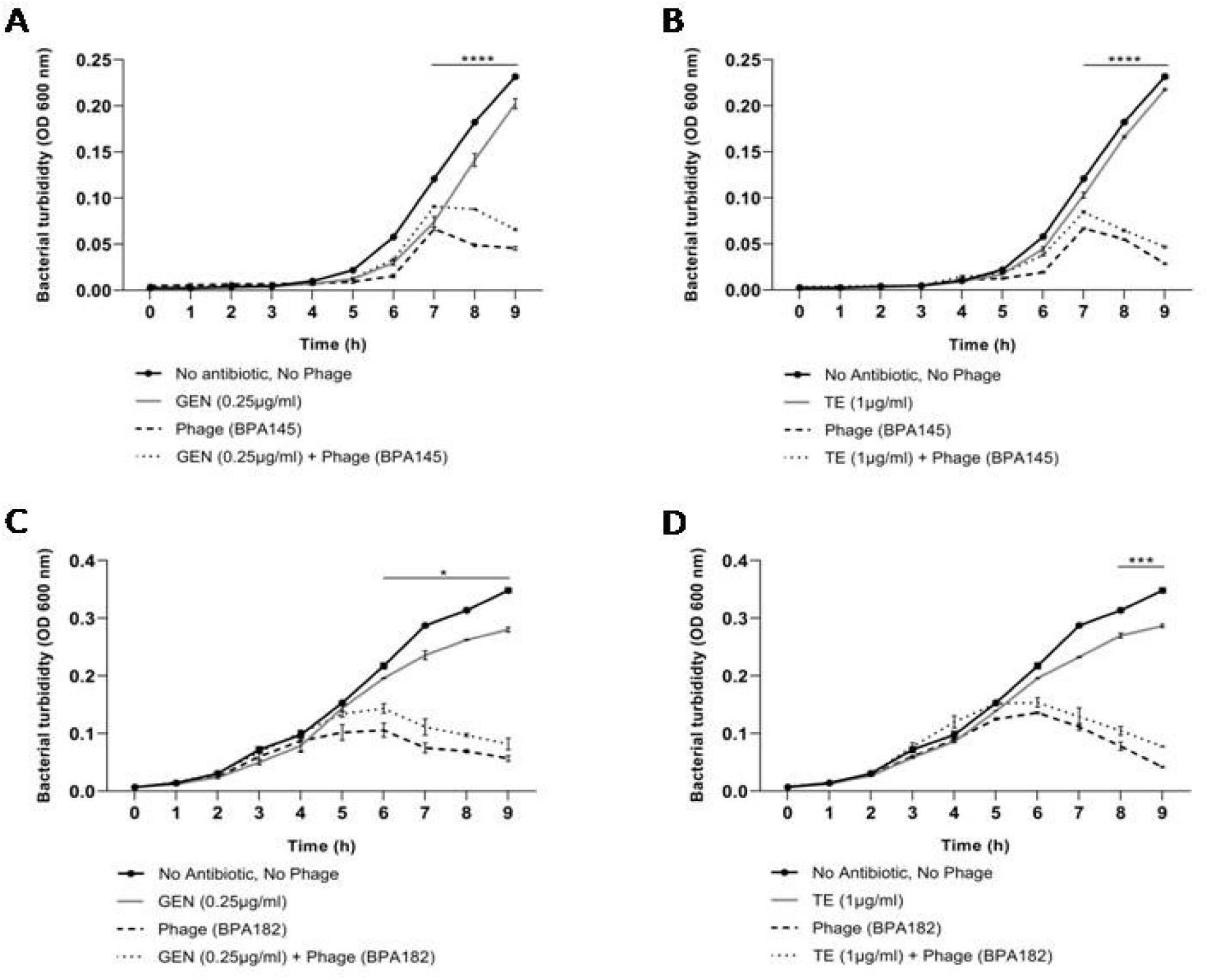
Liquid infection assay: Effect of combination of bacteriophage BPA145 and antibiotics on *A. baumannii* (VTCCBAA1521) (A) GEN (0.25μg/ml) (B) TE (1μg/ml). Effect of combination of bacteriophage BPA182 and antibiotics on *A. baumannii* (VTCCBAA1084) (C) GEN (0.25μg/ml) (D) TE (1μg/ml). BPA145 and BPA182 were used at an MOI of 0.001 and GEN & TE were used at ¼^th^ MIC. Statistical analysis was performed using two way factorial ANOVA with Tukey's test to determine the antagonistic effect (p<0.01) in GraphPad Prism 8.0.2. Error bars depict standard error of mean. Significance levels are depicted as **** p<0.0001; *** p<0.001; * p<0.05. Abbreviations used as gentamicin (GEN), tetracycline (TE).

The evidence gathered through these findings is suggestive of the fact that the antibiotics like aminoglycosides, tetracycline and macrolides which interfere with the ribosomal machinery (8,9,10) of the bacteria makes ribosomes partially inaccessible for use by bacteriophages leading to reduction in their lytic activity. In other terms, the mentioned antibiotics potentially show partial competitive inhibition of bacteriophages and alter their pharmacodynamic potential. This highlights that before considering the combination therapy of bacteriophages with antibiotics for treatment, we should screen them by simple agar overlay in vitro test for synergy / antagonism to rule out any negative impact of treatment. The findings from the study have significant implication in decision making for devising efficacious treatment regimen involving combination of phages and antibiotics to successfully treat multi drug resistant infections in humans/livestock.

## Data availability

All the bacterial and bacteriophage isolates used in this study are submitted in Bacteriophage Repository, National Centre for Veterinary Type Cultures, ICAR - National Research Centre on Equines, Hisar, Haryana, India.

## Acknowledgments

The financial support from Indian Council of Agricultural Research, New-Delhi, India (grant IDOXX04448) is duly acknowledged.

## References

1. Callaway TR, Lillehoj H, Chuanchuen R, Gay CG. 2021. Alternatives to Antibiotics: A Symposium on the Challenges and Solutions for Animal Health and Production. Antibiotics (Basel) 10:471.

2. Międzybrodzki R, Hoyle N, Zhvaniya F, Łusiak-Szelachowska M, Weber-Dąbrowska B, Łobocka M, Borysowski J, Alavidze Z, Kutter E, Górski A, Gogokhia L. 2021. Current Updates from the Long-Standing Phage Research Centers in Georgia, Poland, and Russia, p. 921–951. *In* Harper, DR, Abedon, ST, Burrowes, BH, McConville, ML (eds.), Bacteriophages: Biology, Technology, Therapy. Springer International Publishing, Cham.

3. Comeau AM, Tétart F, Trojet SN, Prère M-F, Krisch HM. 2007. Phage-Antibiotic Synergy (PAS): beta-lactam and quinolone antibiotics stimulate virulent phage growth. PLoS One 2:e799.

4. Schooley RT, Biswas B, Gill JJ, Hernandez-Morales A, Lancaster J, Lessor L, Barr JJ, Reed SL, Rohwer F, Benler S, Segall AM, Taplitz R, Smith DM, Kerr K, Kumaraswamy M, Nizet V, Lin L, McCauley MD, Strathdee SA, Benson CA, Pope RK, Leroux BM, Picel AC, Mateczun AJ, Cilwa KE, Regeimbal JM, Estrella LA, Wolfe DM, Henry MS, Quinones J, Salka S, Bishop-Lilly KA, Young R, Hamilton T. 2017. Development and Use of Personalized Bacteriophage-Based Therapeutic Cocktails To Treat a Patient with a Disseminated Resistant Acinetobacter baumannii Infection. Antimicrob Agents and Chemother 61:e00954–17.

5. Rostkowska OM, Międzybrodzki R, Miszewska‐Szyszkowska D, Górski A, Durlik M. 2021. Treatment of recurrent urinary tract infections in a 60‐year‐old kidney transplant recipient. The use of phage therapy. Transpl Infecti Dis 23:e13391.

6. Furfaro LL, Payne MS, Chang BJ. 2018. Bacteriophage Therapy: Clinical Trials and Regulatory Hurdles. Front Cell Infect Microbiol 8:376.

7. Morris FC, Dexter C, Kostoulias X, Uddin MI, Peleg AY. 2019. The Mechanisms of Disease Caused by Acinetobacter baumannii. Front Microbiol 10:1601.

8. Kotra LP, Haddad J, Mobashery S. 2000. Aminoglycosides: perspectives on mechanisms of action and resistance and strategies to counter resistance. Antimicrob Agents Chemother 44:3249–3256.

9. Nelson ML, Levy SB. 2011. The history of the tetracyclines. Ann N Y Acad Sci 1241:17–32.

10. Vázquez-Laslop N, Mankin AS. 2018. How Macrolide Antibiotics Work. Trends Biochem Sci 43:668–684.

